# Dynamics of sensorimotor reweighting: How light touch alters vestibular-evoked balance responses

**DOI:** 10.1101/2024.04.12.589029

**Authors:** Megan H. Goar, Michael Barnett-Cowan, Brian C. Horslen

## Abstract

Integrated multisensory feedback plays a crucial role in balance control. Minimal fingertip contact with a surface (light-touch), reduces center of pressure (CoP) by adding sensory information about postural orientation and balance state. Electrical vestibular stimulation (EVS) can increase sway by adding erroneous vestibular cues. This juxtaposition of conflicting sensory cues can be exploited to explore the dynamics of sensorimotor reweighting. We used continuous stochastic EVS (0-25Hz; ±4mA; 200-300s) to evoke balance responses in CoP (Exp-1, Exp-2) and segment accelerations (Exp-2). Systems analyses (coherence, gain) quantified coupling and size of balance responses to EVS. We had participants either touch (TOUCH; <2N) or not touch (NO-TOUCH) a load cell during EVS (Exp-1, Exp-2), or we intermittently removed the touch surface (Exp-2) to measure the effects of light touch on vestibular-evoked balance responses. We hypothesized that coherence and gain between EVS and CoP would decrease, consistent with the CNS down-weighting vestibular cues that conflict with light touch. Light touch reduced CoP displacement, but increased variation in the CoP signal explained by EVS input. Significant coherence between EVS and CoP was observed up to ∼30Hz in both conditions but was significantly greater in the TOUCH condition from 12-28.5-Hz. Conversely, EVS-CoP gain was 63% lower in TOUCH, compared to NO-TOUCH. Our findings show that light touch can re-weight vestibular-evoked responses by reducing their size but also increasing high frequency vestibular contributions for sway. This suggests that the CNS can use novel sensory inputs to alter balance behavior but cannot completely ignore a salient balance cue.

**New and Noteworthy:** This study reveals that minimal fingertip contact (light touch) during balance tasks not only diminishes the impact of electrical vestibular stimulation (EVS) on sway, but also enhances the central nervous systems ability to integrate high-frequency vestibular cues. Specifically, light touch decreases the magnitude of EVS-induced sway while increasing coherence with EVS at higher frequencies, illustrating the central nervous system’s capacity to adaptively reweight sensory inputs for improved balance control without fully disregarding dominant cues.

## Introduction

To maintain balance and posture, the central nervous system (CNS) integrates sensory signals by forming a weighted sum of inputs from various modalities (1). This means that some sensory modalities can contribute more to the representation of balance and posture, being weighted heavier, while others contribute less. These weights are typically not equal between modalities (1–4), and the CNS appears able to change these weights (termed ‘reweighting’), depending on the task or environmental conditions (5–9).

Sometimes sensory modalities conflict with each other, presenting contradicting balance relevant cues (1, 3, 6, 7, 10). This can happen when cues that are decoupled from how the body is positioned with respect to gravity from one modality contradict other balance relevant cues. Decoupling sensory cues from the gravitoinertial plane can be caused by sensory dysfunction such as unilateral vestibulopathies (11), certain behavioral contexts such as standing on a boat (12) or virtual reality environments that provoke sickness (13–15). Sensory conflict is probed experimentally with artificial sensory stimulation, such as tendon vibration (16, 17), changing visual objects on a surround projection, or vestibular stimulation (10, 15). Sensory conflicts can perturb postural control, sometimes causing instability and falls (7, 18–20). To help resolve this, the CNS is thought to increase the weight of cues that reliably indicate how the body is positioned with respect to gravity, termed ‘upweighting,’ while decreasing the weight of the cues that do not or are less reliable, termed ‘downweighting’ (1, 6–9). It is not fully known if there are rules for how the CNS determines which cues are indicative of how the body is positioned with respect to gravity, and which are not. However, Bayesian models that form statistically optimal predictions for sensorimotor control based on weighted sensory sums have proven to be effective (21–23). In this study, we assess the dynamics of the reweighting process itself during balance and postural control by using a novel paradigm to assess the effect of introducing light-touch when subjected to noisy vestibular stimulation.

Electrical vestibular stimulation (EVS) can be used to probe the dynamics of vestibulo-motor re-weighting in standing balance and postural control. EVS non-invasively targets the vestibular organs, which are in the inner ear and provide balance relevant feedback about movement of the head (e.g., rotation and translation) and tilt relative to gravity (24). The activation pattern induced by EVS differs from that of physical motion, as it simultaneously stimulates vestibular afferents with various directional sensitivities. Consequently, when EVS signals are sufficiently potent (e.g., >1mA; (25)), the artificial EVS signals can disrupt balance and postural control and decouple evoked balance responses from actual head orientation or movement within the gravitoinertial plane (10, 15). The EVS signal can be manipulated and shaped to the needs of the experimental condition. Noisy (e.g., stochastic or multifrequency) EVS has been used in conjunction with linear systems analyses to assess sensorimotor reweighting of vestibulo-motor balance responses within or between behavioral tasks, reflecting central vestibular reflex modulation (5, 8, 26–31). When responses are more correlated to vestibular input and/or increased in amplitude, it is thought that vestibular weighting on balance or postural control is increased by the CNS.

Other sensory inputs that can be experimentally controlled such as light touch (<2 N) cues from a finger contacting an earth mounted surface, can reduce sway (33). Light touch indicates balance state in an ergocentric frame by coding for body movement with respect to the stable surface (32, 33). In contrast to extending the base of support like when using a cane or walker (34, 35), pressure and shear signals from the fingertips can be combined with other somatosensory cues representing hand, arm, and body posture to inform the CNS of subtle changes in body motion and/or orientation relative to the non-moving, reference surface (32, 33, 36). Since pressure is kept to less than 2 N, it is thought that light touch does not provide enough mechanical force to stabilize the body (37) but reduces postural sway and/or centre of pressure (CoP) displacements through sensory stabilization (36–38).

There is an opportunity to use continuous EVS and light touch to determine how the CNS responds to sensory conflict. When both cues are available, the CNS must determine which sensory cues are noisy or disruptive to balance and posture and which cues are veridical or able to best stabilize the body. Assuming the behavioral goal is to stand still, we would expect the CNS to down-weight the noisy cue (EVS-driven vestibular cues) and up-weight the noise-free cue (light touch) for sensorimotor control of balance and posture. We can test this assumption by measuring the vestibular systems’ influence on CoP displacement both in the presence and absence of light touch to uncover the extent to which the CNS modulates vestibular-motor control of posture and balance in response to sensory conflict.

Systematically manipulating postural control via two unique sensory inputs allows for an experimental approach to assess the CNS’s ability to integrate a novel balance relevant sensory input and modulate a conflicting one. While we understand sensory reweighting during conflict occurs, we do not know how long it takes for one balance relevant sensory modality to modulate another. It takes approximately 150 ms for the earliest responses to EVS to be reflected in ground reaction forces, with the largest responses occurring 300-350 ms after onset of EVS (5, 39). The perceived timing of EVS occurs slightly later, around 438 ms (41, 42). Conversely, correlated changes in CoP lag continuous and changing light touch cues by 275-300 ms (40). However, Sozzi and colleagues (38) showed that it can take 0.7-2 s for newly introduced passive light touch cues to lead to changes in CoP, but only up to 1 s for removal of light touch to affect CoP. The timing of vestibulo-motor adaptations has also been measured in response to changes in balance task. Vestibulo-muscular coupling begins to decrease 150-200 ms after switching from active balancing to externally stabilized balancing (8) and 1.5 s after introducing sensorimotor delays to active balancing using the same paradigm (9). Furthermore, there is evidence of anticipatory vestibulo-motor adaptation with voluntary changes in balance behavior. Tisserand and colleagues (31) found that EVS-CoP coherence decreased 259-435 ms before starting walking, and during the last step before stopping walking. Despite the body of literature focusing on timing of responses to changes in vestibular or light touch cues, we do not know how long it takes to resolve sensory incongruency between cutaneous and vestibular modalities. Longer latencies for the CNS to respond may vary depending on how long it takes to determine which cues are more reflective of how the body is positioned with respect to gravity (43). Taking these studies together, timing of vestibulo-motor adaptations can range from reactionary to anticipatory and may be asymmetric upon introduction or removal of cues, perhaps depending on how difficult or complex the adaptations are for the CNS to execute.

The purpose of this study was to assess the manner and duration in which light touch sensory cues influence vestibulo-motor control of posture and balance. In our first experiment, we used stochastic EVS to evoke balance responses either in the presence or absence of light touch to determine how the CNS changes vestibulo-motor coupling when additional conflicting sensory information becomes available. We hypothesized that vestibulo-motor coupling, as reflected by EVS-CoP coherence and gain, would decrease when light touch was available. In our second experiment, we used regular, intermittent, transitions between presence and absence of light touch to determine the time course of EVS-CoP and EVS-segment acceleration coherence and gain adaptations in response to an additional sensory cue. While CoP adaptations to light touch have been shown to occur within 0.7-2 s of touch being introduced (38), vestibulo-motor reweighting can occur as fast as 150 ms after a change in sensory conditions (8). Thus, we hypothesized that EVS-CoP coherence and gain would begin to decrease or increase approximately 150 ms after introduction or removal of light touch, respectively.

## Methods

### Participants and ethical approval

A sample of 16 participants (n females = 8) was chosen for experiment 1 and a sample of 10 participants (n females = 7) was chosen for experiment 2. For both experiments, neurotypical young adults (18-35 yrs. old) with no self-reported neurological or orthopedic issues that affect standing balance control were recruited. All participants gave written informed consent before their participation in the experiments, and all methods were reviewed and approved by the University of Waterloo Research Ethics Board (REB #44217).

### Experimental Protocol

Participants stood barefoot on 2 force plates (AMTI OR6-5, Advanced Mechanical Technology, Inc., Watertown, MA, USA), and their foot length was measured. The feet were then positioned such that each foot was on a different plate and the lateral edges of their 5th metatarsals were spaced equal to their foot length (5). Foot position was then marked to ensure consistent placement between trials. An experimenter monitored foot position during and between trials.

Head position was controlled to both align responses to EVS in the anterior-posterior (AP) or medial-lateral (ML) direction and to keep these responses in the same direction across participants for each trial and both experiments. Head angle was set for all trials at approximately 18 degrees above Reid’s plane, as this head orientation aligned responses to the frontal-sagittal plane (25). Responses were aligned in the ML direction when the head was forwards and responses were aligned in the AP direction when head was to the side. For AP trials, the perceived vestibular perturbation was aligned with the feet. Participants were instructed to maintain ML or AP head posture throughout each trial duration. Responses in the AP direction were shown to have greater effect sizes than responses in the ML direction during experiment 1, which is why ML trials were not conducted in experiment 2 and only results from AP trials were used and reported (see supplemental material Fig. 1 for ML results).

**Figure 1:**
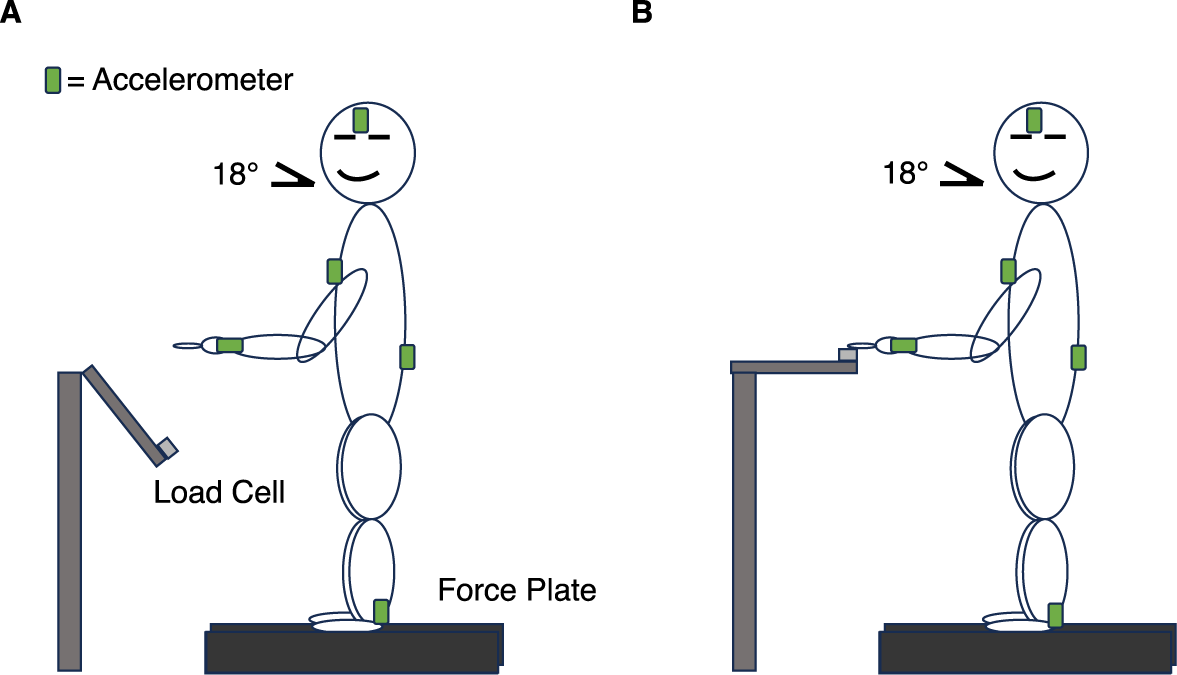
Setup for NO-TOUCH (A) and TOUCH (B) Conditions. Participants, with eyes closed and head turned 90 degrees yaw and tilted 18 degrees above Reid’s plane, stood still on two force plates (black rectangles) to record CoP displacement and wore accelerometers (green rectangles) for measuring segmental acceleration. In TOUCH (B), the load cell is present; it is withdrawn in NO-TOUCH (A).

Accelerometers (Shimmer3 IMU, Shimmer, Dublin, Ireland) were worn at the head, sternum, low back, dominant wrist, and ankle for experiment 2 only. They were secured the same way every time, using appropriate landmarks on the body (center of the frontal region of the head, sternum, low back, wrist joint line, distal Achilles tendon between malleoli) and accelerometer itself. The accelerometers were aligned and secured such that the unit positive Z and Y axes were aligned in the anatomical anterior and superior directions, respectively. The accelerometers were fastened with bands that had holders for the accelerometers, except for the head, which was secured with a tensor headband. The positioning of the accelerometers on the participant is illustrated in figure 1.

Figure 1 also illustrates the two conditions used for both experiments: NO-TOUCH and TOUCH. Prior to collection, a load cell was calibrated so that a visual display of the load cell signal in volts could be interpreted in Newtons. An experimenter monitored light touch vertical force during both the TOUCH (1-2 N) and NO-TOUCH (0 N) conditions on a computer screen. Experimenters could give verbal instructions to participants to adjust their load as needed. Participants were asked to slightly flex their dominant arm (left hand was used for 1 participant of experiment 1) at the shoulder and elbow so that their index finger rested around hip height in a comfortable position, slightly in front of the body. For experiment 1, a 1 cm diameter circular load cell (Model SLB-100 compression load button with 445 N range, Transducer Techniques LLC, Temecula, CA, United States; amplifier: Model 3270 strain gage bridge conditioner, Daytronic dba Dranetz Technologies, Edison, NJ, USA) was permanently fixed in position for the duration of the experiment. For experiment 2, the load cell (burster 8523-5020-N000S000 tension-compression load cell with 20 N range: A-TECH Instruments LTD, Scarborough, ON, Canada; amplifier: Model 3270 strain gage bridge conditioner, Daytronic dba Dranetz Technologies, Edison, NJ, USA) was embedded within a 6 x 6 cm plastic square to increase surface area and positioned so that it could be raised by the experimenter using a lever system into a locked position for participants during TOUCH and lowered for participants during NO-TOUCH. Participants were instructed to maintain their arm posture while an experimenter controlled transitions between NO-TOUCH and TOUCH conditions (Fig. 1).

A novel device was custom-built to suddenly switch the participant between NO-TOUCH and TOUCH conditions for experiment 2. In the NO-TOUCH condition, an experimenter pushed the lever arm holding the load cell down, disengaging magnets holding it in place, and causing the load cell arm to swing downwards and away from the participant (Fig. 1A). In the TOUCH condition, an experimenter brought the arm upwards, reengaging the magnets, and the load cell arm was held in place under the participant’s index fingertip (Fig. 1B).

For the TOUCH condition, the participant was instructed to use their index finger flexors controlling the metacarpophalangeal joint to isometrically press the load cell to a minimum vertical load of 1 N and maximum of 2 N (33, 37). For the NO-TOUCH condition, the participant was instructed to use their index finger extensors controlling the metacarpophalangeal joint to isometrically hold the index finger against gravity, above where they expected the load cell to be (experiment 1), or the experimenter removed the load cell and participants held their finger where the load cell would be if in the fixed TOUCH position (experiment 2). Participants were instructed to maintain similar arm postures for the NO-TOUCH and TOUCH conditions for both experiments so that this posture did not contribute to changes in the relationship between vestibular inputs and balance or postural control.

### EVS

For both experiments, bipolar binaural EVS was delivered percutaneously above the mastoid processes to stimulate the nearby vestibular afferents. The anode was worn on the right and cathode on the left. A stochastic signal was generated for each trial using a custom LabVIEW code (National Instruments Corp., Austin, Tx, USA), which was digitally low pass filtered at 25 Hz (5, 26, 29). The digital signal was then converted to analog and fed to a stimulator (STMISOLA Linear Isolated Stimulator, Biopac Systems, Inc., Goleta, CA, USA). The EVS peak amplitude was 4.5 mA with a 0 mean.

### Sampling

All signals were sampled at 1024 Hz. EVS, force plate ground reaction forces and moments, and load cell were sampled at 1024 Hz using a custom LabVIEW script (Data Acquisition Board: National Instruments Corp., Austin, Tx, USA; LabVIEW: National Instruments Corp.). Accelerometry data were sampled at 1024 Hz using ConsensysPRO software (Consensys, Brooklyn, NY, USA) and the Shimmer3 data acquisition board. The accelerometry data were synchronized in time with EVS and force plate data using a TTL synch pulse delivered simultaneously to both collection systems at the beginning of each trial.

### Trial Structure

Before data collection began, participants received a short exposure to the EVS for 10 seconds. This was so participants could understand how stimulation feels, report any discomfort, and allow the experimenter to adjust electrode placement or increase electrode gel if stimulation was uncomfortable, before the recorded trials. Participants were permitted multiple exposures before starting data collection if they wished. Once trials began, participants were stimulated for the entire duration of the trial.

Prior to any trial type, participants were asked to close their eyes and stand as still as possible until an experimenter informed them that the trial was complete. This was done to limit sensory input from the visual and proprioceptive systems. Participants took seated breaks of at least 1 min between each trial.

There were 3 trial types: baseline, single state and switching. Single state and baseline trials were the same and were reported for both experiments. Switching trials were only collected in experiment 2. Only EVS and CoP data (no segmental acceleration) from switching trials are reported from experiment 2.

The 2 baseline trials were 60 s in duration, and participants were instructed to either not touch (NO-TOUCH) or touch (TOUCH) the load cell for the whole duration. There was no EVS during these trials, as these data were used to characterize participants’ responsivity to the light touch independent from EVS. Two single state trials were conducted, which were 200 s and trials were again either NO-TOUCH or TOUCH, but with simultaneous EVS the whole time. These data were used to address the relationship between EVS and CoP and how this was modulated by light touch.

For switching trials (experiment 2 only), a passive paradigm was used, where the experimenter controlled a lever system that raised the load cell to a fixed position touching the index finger or withdrew the load cell. The switching trials were 300 s and participants regularly switched between TOUCH and NO-TOUCH conditions every 6-15 s with simultaneous EVS. The intervals between transitions were randomized to prevent prediction of the switching events. These trials were used to see how the EVS and CoP relationship changes over the transitions to NO-TOUCH or TOUCH.

### Outcome Measures

Center of pressure (CoP) was calculated offline using a custom MATLAB script (R2021b, MathWorks Natick, MA, USA) from ground reaction forces and moments from the 2 force plates. CoP data were low pass filtered at 50 Hz (dual pass) with an effective 4^th^ order Butterworth filter. Such a high low-pass cutoff frequency was chosen to ensure the highest EVS stimulation frequencies (i.e., 25 Hz and above) were not removed from the CoP signal prior to analyses.

Baseline-corrected root mean square (RMS) of CoP was used to assess the variance in CoP displacement amplitude for the no-EVS baseline and single-state EVS trials in both experiments. These analyses permitted assessment of the effects of light touch on CoP displacement independent from EVS. CoP data were baseline corrected by subtracting the mean CoP calculated from the entire trial duration (60 s for baseline, 200 s for single state). CoP RMS was then calculated from the baseline corrected data. All calculations were performed using a custom MATLAB script.

Accelerometry data were filtered offline with a 4^th^ order Butterworth low-pass filter with a cutoff frequency of 50 Hz (dual pass) using a custom MATLAB script. Accelerations in the dimensions corresponding with the expected EVS-evoked body sway at each segment were used for further analyses (i.e., AP for the sternum and dominant arm, and ML for the head).

Accelerometer sampling occasionally dropped due to wireless connectivity issues. To fill in drops in the data of gaps less than 0.2 s, a cubic spline was used in the custom script (44). Since cubic splines will give large artifacts for gaps greater than 0.2 s, linear interpolation was used to fill these larger gaps. This makes the final set have the desired 1024 Hz sampling rate, thereby facilitating comparison with other data from this experiment.

Load cell data representing light touch force were low pass filtered at 5 Hz (dual pass) with a 4^th^ order Butterworth using a custom MATLAB script. The load cell data were used to determine when a participant was touching the stable reference. Participants were in the TOUCH condition when the recorded force was 3 standard deviations above the unloaded level. An unloaded baseline was established by calculating the mean and standard deviation of the recorded signal when the load cell was unloaded. This baseline reflects both any effects of gravity on the load cell, and signal noise in the recorded system.

For switching trials, EVS and CoP were sectioned around a transition marked in the load cell data (i.e., either transition from NO-TOUCH to TOUCH, or vice versa). Each section of data was 12 s long, with 6 s on either side of the transition. Due to randomization of transition timing, not all participants achieved the same number of transitions either from one condition to the other, or as other participants in the same transition direction. We, therefore, excluded data from some participants to ensure that all participants contributed an equal number of transitions to the final analysis. This was done by matching the participant with the fewest transitions. In total, there 380 transitions from TOUCH to NO-TOUCH (38/participant), and 370 transitions from NO-TOUCH to TOUCH (37/participant).

### Linear Systems Analyses

A linear systems approach was used to determine the relationship between EVS and CoP or linear acceleration in the frequency and time-frequency domains. Coherence and gain estimates between the EVS and CoP were calculated using the NeuroSpec 2.0 code. NeuroSpec 2.0 (www.neurospec.org) is a freely available archive of MATLAB code intended for statistical signal processing and based on the methods of Halliday and colleagues (45) and is well esablished in the literature (5, 8, 9, 26, 27, 30).

Coherence was used to express the amount of variation in the output signal that can be explained by the input signal across 0-30 Hz. It is a measure of correlation bounded between 0 and 1 for each frequency bin and indicates where in the frequency spectrum signals are correlated. Zero represents the case of independence, while 1 represents the case of a perfect linear relationship. It is calculated using equation 1, where the cross spectrum of the input and output is divided by the input and output auto-spectra (x is input signal, i.e., EVS; y is output signal, i.e., CoP) For our purposes, we defined EVS as the input signal and our dependent measures (CoP, linear acceleration) as the output signal.

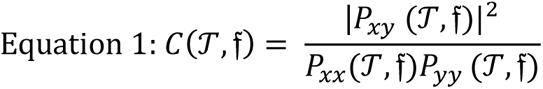

For all coherence calculations, baseline corrected CoP was amplitude normalized by dividing the baseline corrected CoP by the maximum CoP amplitude. This was to ensure that changes in coherence were due to changes in the cross spectrum between EVS and CoP, and not changes to their respective auto-spectra. CoP amplitude normalization and its effect on CoP power can be seen in supplementary materials figure 2.

**Figure 2:**
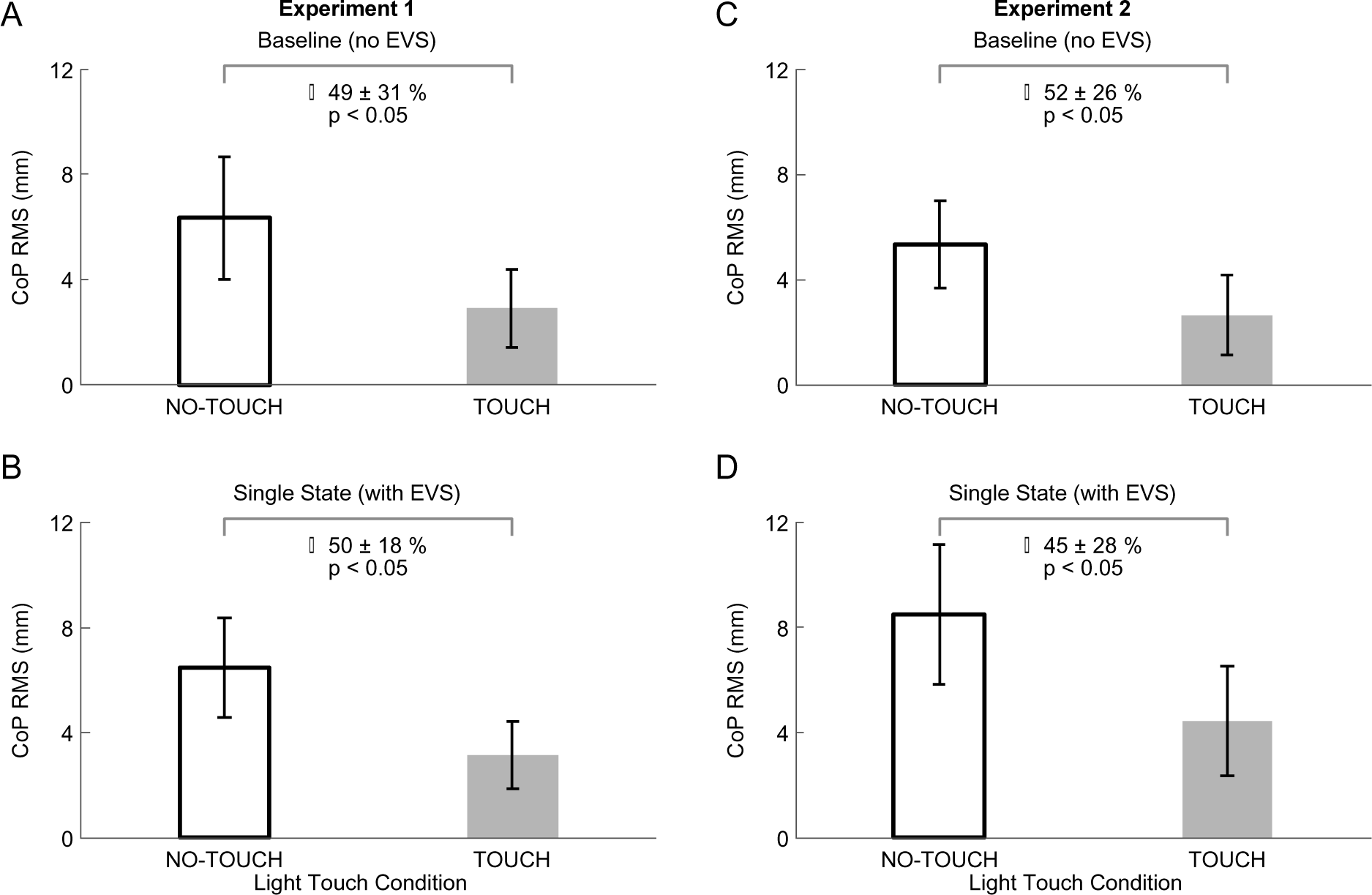
CoP RMS Differences Between NO-TOUCH and TOUCH. Averages for baseline (EVS; A, B) and single state (no EVS; C, D) across experiments. Data averaged from 15 participants (experiment 1) and 10 participants (experiment 2), with NO-TOUCH (black outline) and TOUCH (grey). Error bars indicate one standard deviation.

Gain reflects the scale relationship between input and output signals; in this study it reflects the amplitude of the CoP displacement to a given EVS current. Interpreting gain is conditional on significant coherence between signals, so any frequencies where coherence was not significant were excluded from the gain analyses. It is calculated by using equation 2, where the magnitude of cross-spectrum between an input and output is divided by the input auto-spectrum.

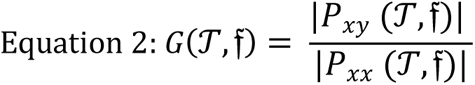

Single-state coherence and gain were calculated for each participant. Participants contributed 200 s of EVS and CoP (or segmental acceleration; 100 non-overlapping 2s bins; 2048 samples/bin), beginning at the onset of EVS, for each of the TOUCH and NO-TOUCH conditions. Coherence and gain between EVS and CoP measures were calculated with a 0.5 Hz resolution from 0.5 Hz to 30 Hz for each participant. A 2 s window (5 Hz resolution) was needed to resolve frequencies at 0.5 Hz, where most CoP power occurs (49). The frequency range was selected because vestibulo-motor coherence has been found up to 30 Hz in previous studies (5, 9, 26, 27, 30, 30).

Group-wide analyses of single-state coherence and gain were calculated from single data arrays containing equal amounts of data from each participant (same number of bins as individual participant analysis). Data were concatenated into arrays such that input and output data were matched sequentially (i.e., EVS data from participant 1 aligned with CoP data from participant 1). This led to a final sample of 1500-2s bins from experiment 1 (n=15 participants) and 1000-2s bins from experiment 2 (n=10).

Within-subjects analyses of coherence were used to confirm that group-wide differences persisted after controlling for inter-participant differences. Cumulative sums of EVS-CoP coherence for individual participants (i.e., without pooling) were calculated across a high frequency bandwidth (13-30 Hz) using custom MATLAB scripts. This range was chosen based on observations made from pilot testing. Cumulative sums were calculated by adding the coherence at each frequency. Average and standard deviation of the participant cumulative sums were compared for each condition.

Switching trial data were concatenated into single EVS and CoP arrays (see single state, above). Coherence and gain were calculated using sliding 2 s windows, with 0.01s step time, (2048 samples, 0.5 Hz resolution; (8, 46)). This led to 1201 total overlapping bins, with 600 on each side. To determine the mean time to for coherence and gain to change around a transition to NO-TOUCH or TOUCH, coherence and gain were averaged across frequencies where there were significant differences found across conditions for single state analysis (13-25 Hz for coherence, 0.5-5 Hz for gain) at each time step. Latency of change in either coherence or gain was calculated in reference to the original state (i.e., if transition is from NO-TOUCH to TOUCH, then in reference to NO-TOUCH). A change in coherence or gain was taken to occur when the time-dependent average signal either rose above or below a mean ±2 SD threshold calculated from the first 2 s of the data.

A separate analysis was conducted on the switching trial data to confirm that the switching trials data showed comparable results to the single-state trials. We separated the switching trial data into post hoc NO-TOUCH and TOUCH arrays and conducted single-state analyses prior to time-dependent ones. Eight seconds on either side of the 380 transitions to NO-TOUCH were used. This led to 1520 bins per NO-TOUCH (after transition) and TOUCH (before transition) conditions (153 bins per participant; see supplementary materials Fig. 3).

**Figure 3:**
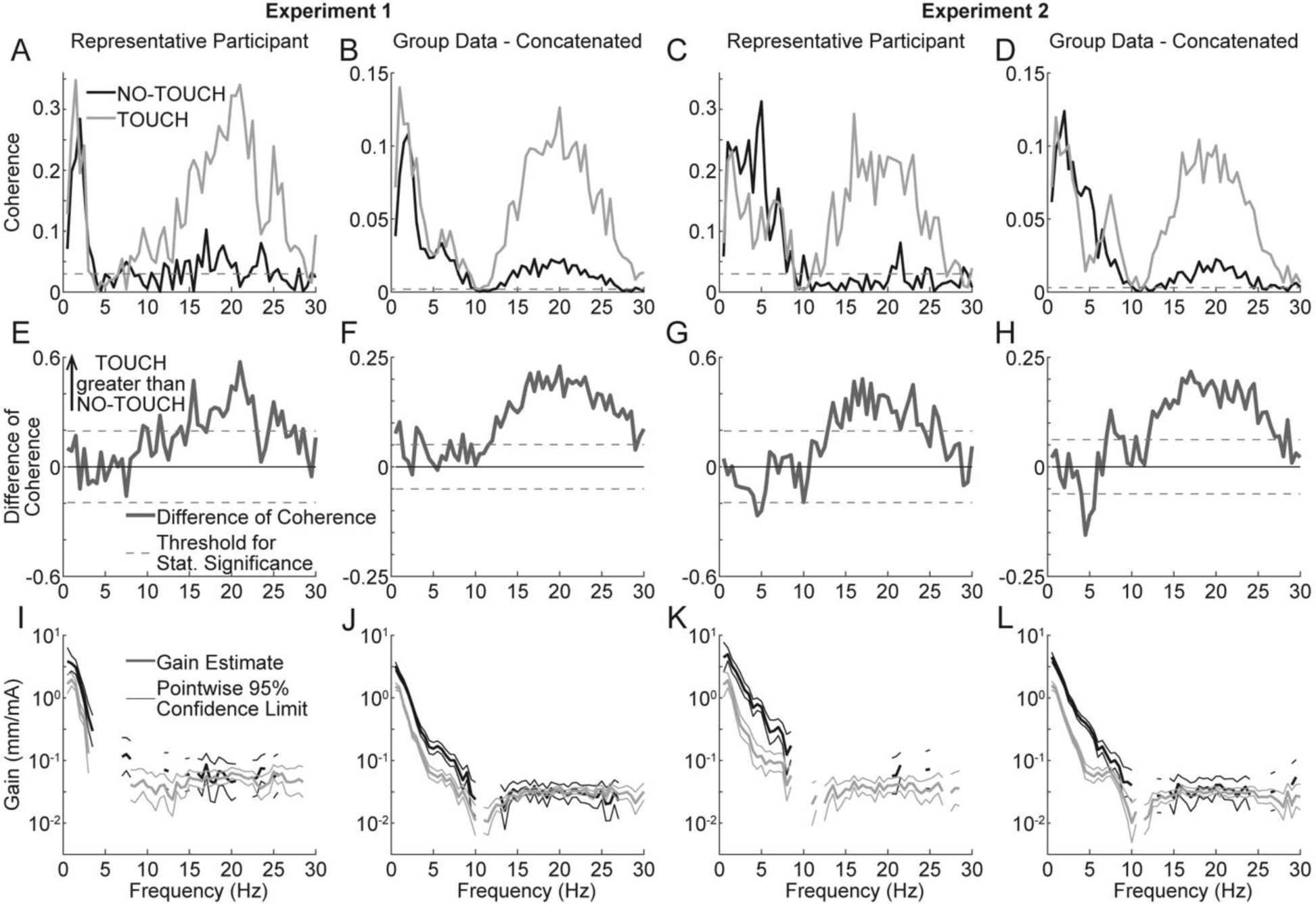
Coherence and Gain Differences between TOUCH and NO-TOUCH. Panels A-D illustrate EVS-CoP coherence with NO-TOUCH (black) and TOUCH (grey) for a representative participant from experiment 1 (A), pooled data from experiment 1 (B), a representative participant from experiment 2 (C), and pooled data from experiment 2 (D). A dashed horizontal line indicates the 95% confidence limit threshold for significant coherence. Panels E-H show difference of coherence (black solid lines) with positive/negative 95% confidence limits (black dashed lines), where positive values indicate frequency ranges where TOUCH EVS-CoP coherence is greater than NO-TOUCH coherence. Panels I-L present EVS-CoP gain on a log scale with pointwise 95% confidence limits (thin lines), excluding frequencies without significant coherence for clarity. Statistically significant differences in gain can be seen where 95% confidence limits do not overlap.

In post-processing we revised bin width to a 4 Hz resolution with window that were 0.25 s long (256 samples) to increase the time resolution and better resolve the high frequency coherence changes over time, using the smallest time window that can be used (see time dependent analyses in results). However, reducing the frequency bins also reduces total coherence amplitude, increases variability in the signal, and increases the prevalence of any signal drift. Lower frequencies also cannot be resolved, so these processing parameters were used to focus on frequencies above 12 Hz only, where high frequency differences between NO-TOUCH and TOUCH were found during single state analysis. Setting thresholds based on 2 SD was not as effective in discerning the differences in coherence levels around the transitions as the original time dependent analysis, so coherence was averaged from a larger range of 12-32 Hz (instead of 13-25 Hz) to increase total coherence amplitude and thereby increase the amplitude of coherence changes around the transition.

### Statistical analyses

A repeated measures t-test or a Wilcoxon-signed rank test, depending on if the assumption of normality was met, was used to see if CoP RMS was significantly different across the NO-TOUCH and TOUCH conditions for baseline and single state trials. Prior to this, a Shapiro-Wilk test confirmed whether data from the single state NO-TOUCH and TOUCH conditions were normally distributed, along with the baseline NO-TOUCH and TOUCH conditions. P < 0.05 was interpreted as a failure to meet the assumption of normality. Mean CoP RMS for the conditions, mean percentage change from NO-TOUCH to TOUCH, and standard deviations of percent change were also calculated.

A difference of CoP spectra statistical test was used to see how TOUCH influenced group concatenated CoP power. This test calculates the log ratio difference of spectra, testing the hypothesis of equal values at each Fourier frequency (45). Standardized differences in spectra were estimated at common frequencies (0.5-30 Hz here) between 2 conditions. Any frequencies where the standardized difference of spectra exceeded the 95% confidence limits (based on frequency distribution) were considered statistically different. Differences in spectra across conditions can be seen in supplementary figures 2I and 2K, and how amplitude normalization changes these differences in figures 2J and 2L.

Mean 95% confidence limits of coherence were calculated across all bins for the individual and group-wide concatenated data. The coherence 95% confidence limit was used as a threshold for significant within-conditions coherence between input and output signals.

Significant within-conditions coherence was a prerequisite for all subsequent analyses. The frequency ranges with significant coherence between NO-TOUCH and TOUCH conditions were recorded and compared.

Individual and group level differences in coherence between NO-TOUCH and TOUCH conditions were examined using a difference of coherence test, based on the methods of Rosenberg and colleagues (48) and Amjad and colleagues (47). This statistic is a modified Χ² test, testing the assumption that the coherence estimates are equal with normally distributed variance across conditions. Standardized differences in coherence were estimated at common frequencies (0.5-30 Hz here) between 2 conditions. Ninety-five % confidence limits were developed based on the Fisher transform (tanh^-1^) of the square root of the coherence values. Any frequencies where the standardized difference of coherence exceeded the 95% confidence limits were considered statistically different.

Paired samples t-test was used to evaluate the within-subjects’ differences of the cumulative sum of coherence for high (13-30 Hz) frequency bandwidths between NO-TOUCH and TOUCH. The Shapiro-Wilk test confirmed the assumption of normality between the samples for NO-TOUCH and TOUCH; P < 0.05 was interpreted as a failure to meet the assumption of normality. Cohen’s d was used to quantify effect sizes of the changes in coherence between conditions. Mean cumulative sum for the conditions, mean percentage change from NO-TOUCH to TOUCH, and standard deviations of percent change were also calculated.

Pointwise 95% confidence limits for gain were calculated for all bins of individual and group concatenated data to determine when there were significantly different gain values across conditions. Non-overlapping regions of the confidence limits at the respective frequencies distinguished when these significant differences occurred (5). When overlapping between the confidence limits occurred, gain values were not considered significantly different. Within the regions where gain was significantly different across conditions, the range and mean of percentage change were recorded to capture effect size.

## Results

### Post Collection Exclusion

Of the sixteen people recruited for experiment 1, one participant dropped out due to time restraints. Of the ten people recruited for experiment 2, one participant did not have any accelerometer data, and data from the head, low back, wrist, and ankle segments each were missing from one participant each, due to wireless connectivity issues. For the baseline trials, one participant in each experiment was excluded due to data logging errors. Thus, 15 participants were included in the final analysis for experiment 1 (14 in baseline), and 10 participants were included in the analyses for experiment 2 (8-9 samples from 9 participants for accelerometry).

### CoP-RMS

Light touch reduced CoP RMS in baseline and single state trials. Non-parametric analyses were used to test for differences in CoP RMS between conditions for all tests except for the single state NO-TOUCH TOUCH comparison in experiment 1 (all Shapiro-Wilk P values < 0.05). There was a statistically significant reduction in CoP RMS from NO-TOUCH to TOUCH that was seen in both experiments for no EVS baseline (exp 1: w_13_= 102, P < 0.001; exp 2: w_8_= 44, P= 0.008) and single state trials (exp 1: t_14_= 7.33, p < 0.001; exp 2: w_9_= 53, P = 0.006). Figure 2 shows average CoP RMS for baseline and single state trials and the average reductions from NO-TOUCH to TOUCH, expressed as a percentage change from NO-TOUCH CoP RMS (exp 1: baseline = 49±31%, with EVS = 45±18%; exp 2: baseline = 52±26%, with EVS = 50±28%; Fig. 2).

### EVS-CoP Relationship

Light touch at the fingertip modulated the relationship between vestibular inputs and balance/postural control. This was observed in both experiments for pooled within-conditions EVS-CoP coherence estimates and, on a participant-by-participant basis (Figs. 3A-D). In both the representative and group-wide data for experiment 1, we observed significant EVS-CoP coherence in the NO-TOUCH condition (Figs. 3A and 3B black lines above grey dashed lines; representative participant: 0.5-3.5 Hz, 7-8 Hz, 16-20.5 Hz, 22.5-24 Hz; group-wide: 0.5-10 Hz, 12.5-27 Hz). Likewise, significant EVS-CoP coherence was found in the TOUCH condition (Figs. 3A and 3B solid grey lines above grey dashed lines; representative participant: 0.5-3 Hz, 8-28.5 Hz, 22.5-24 Hz; group-wide: 0.5-10 Hz, 11-30 Hz). Similar patterns of within-conditions coherence were observed in experiment 2. In experiment 2, there was significant EVS-CoP coherence in the NO-TOUCH condition (Figs. 3C and 3D solid black lines above grey dashed lines; representative participant: 0.5-8.5, 20.5-21.5 Hz; group-wide: 0.5-10 Hz, 14.5-24 Hz). There was also significant EVS-CoP coherence in the TOUCH condition (Figs. 3C and 3D solid grey lines above grey dashed lines; representative participant: 0.5-8.5 Hz, 11-26.5 Hz; group-wide: 0.5-10.5 Hz, 11.5-30 Hz).

Significant differences in coherence between NO-TOUCH and TOUCH conditions were observed at both the single-participant and group-wide levels in experiments 1 and 2 (Figs. 3E-H). Our representative participant from experiment 1 showed significantly greater EVS-CoP coherence between 15-16.5, 17.5-23 and 24-26 Hz in TOUCH, compared to NO-TOUCH (Fig. 3E). At the group level, there was a significant increase in EVS-COP coherence from 12-28.5 Hz in TOUCH, compared to NO-TOUCH (Fig. 3F). In experiment 2, our representative participant (13-24 Hz, Fig. 3G) and group-wide data (7-9 and 12-27.5 Hz, Fig. 3H) had significantly greater coherence in the TOUCH, compared to NO-TOUCH. In sum, light touch increased the variation in the CoP signal that could be explained by EVS, particularly at high frequencies.

The group-wide changes in coherence reflected in the pooled data were also seen when accounting for individual differences. The assumption of normality between for NO-TOUCH and TOUCH samples was confirmed for both experimental data sets (all p > 0.05); There was a large increase across participants in high frequency (13-30 Hz) cumulative sum of coherence during TOUCH, compared to NO-TOUCH. For experiment 1, the average cumulative sum for high frequency NO-TOUCH coherence was 0.79 and for TOUCH was 3.04, which represented a 323 ± 323% increase from NO-TOUCH to TOUCH (Fig. 4B; t_14_ = -5.67, P < 0.001, Cohen’s d = - 4.04). For experiment 2, the average cumulative sum across participants for high frequency NO-TOUCH coherence was 0.74 and for TOUCH was 2.92, representing a 323 ± 194% increase from NO-TOUCH to TOUCH (Fig. 4C; t_9_ = -5.61, P < 0.001, Cohen’s d = -2.41). Increases in coherence during TOUCH persist when accounting for inter-participant differences, with a large effect size.

**Figure 4:**
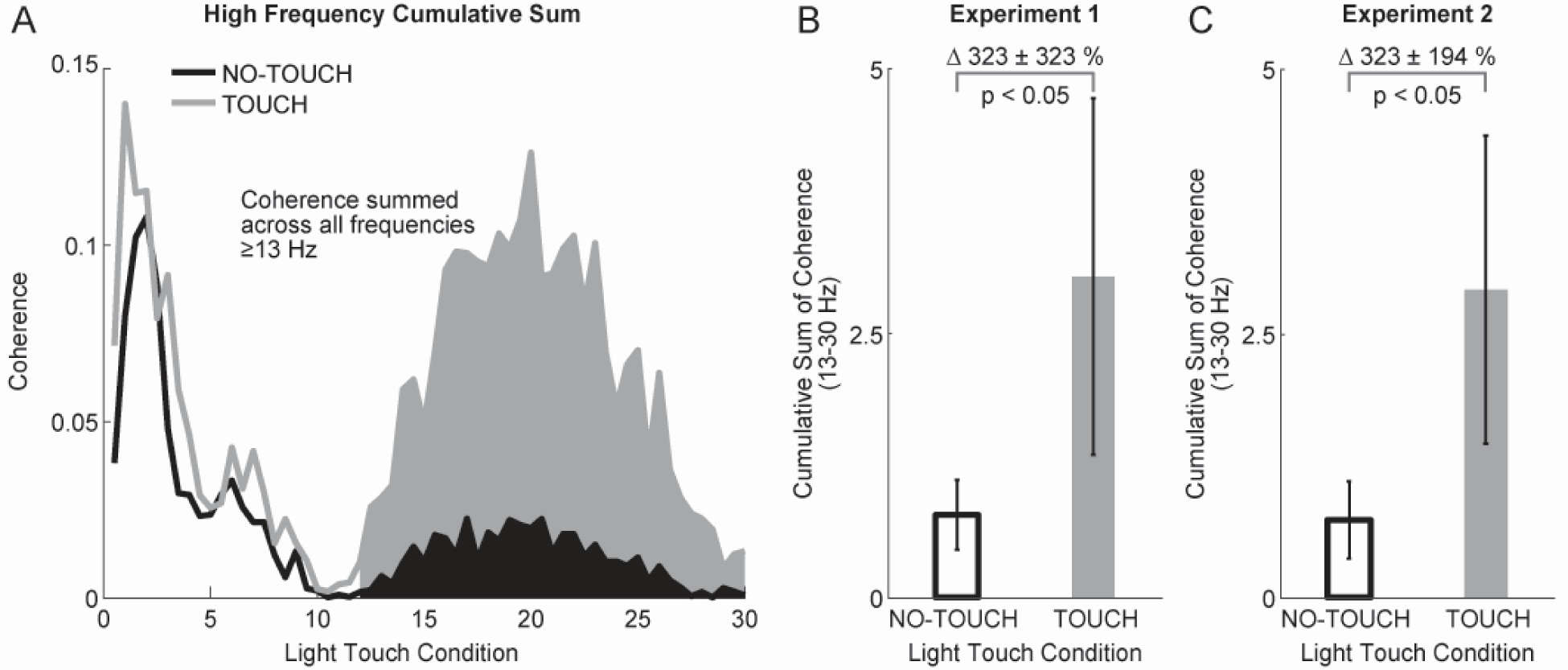
High Frequency Coherence Cumulative Sum Differences Between TOUCH and NO-TOUCH. Panel A illustrates calculation of cumulative sum of EVS-CoP coherence, including frequency range included in the calculation (13-30 Hz, shaded regions). Panels B (experiment 1) and C (experiment 2) compare the average cumulative sum of coherence for NO-TOUCH (black) and TOUCH (grey); where TOUCH cumulative sum was decreased 323% (both experiments 1 and 2), compared to NO-TOUCH. Error bars in B and C represent standard deviation.

Opposite to the direction of effect that light touch had on coherence, EVS-CoP gain was significantly decreased from NO-TOUCH to TOUCH in both experiments at both the single participant and group-wide levels. For the experiment 1 representative participant (Fig. 3I), gain was significantly lower in TOUCH compared to NO-TOUCH from 1-3 Hz. For the pooled data (Fig. 3J), gain was significantly lower in TOUCH compared to NO-TOUCH from 0.5-8 Hz, with a mean reduction of 58% (range of 38-68%). For experiment 2, we observed significantly lower gain in the TOUCH compared to NO-TOUCH condition for the representative participant (1-7.5 Hz, Fig. 3K) and group-wide data (0.5-7 Hz, average 68% reduction, range of 47-83%, Fig. 3L). Significant decreases in TOUCH can be seen where there is complete separation of gain estimates (thick solid lines) and their confidence limits (thin lines; Figs. 3I-L). The gains always peaked at the lowest frequency represented (0.5 Hz) and, on visual inspection, appeared to decay logarithmically as frequency increased for both experiments. These results show that the amplitude of the balance response to a given vestibular input was decreased during TOUCH, meaning there is less CoP displacement per unit of EVS stimulus.

### Modulations of EVS-CoP Relationship Over Time

#### Switching from TOUCH to NO-TOUCH

Since single-state and switching trial data were collected in different trials, we first confirmed that the switching trials data showed similar effects to single-state data when treated in the same way prior to proceeding with the time-dependent analyses. In brief, we observed a significant increase in high frequency coherence, and decrease in gain at low frequencies, in TOUCH, compared to NO-TOUCH in the switching trials data when examined using single-state analyses methods. These data are presented in detail in supplementary materials figure 3.

The time-dependent analysis revealed that coherence decreased, while gain increased, before the load cell was removed from the finger during transitions to NO-TOUCH. Time-dependent coherence is shown in figure 5A, where significant coherence is denoted with a heat map. High frequency coherence (>13 Hz) in teal began to decrease 1.82 seconds before the finger lost contact with the load cell (Fig. 5B).

**Figure 5:**
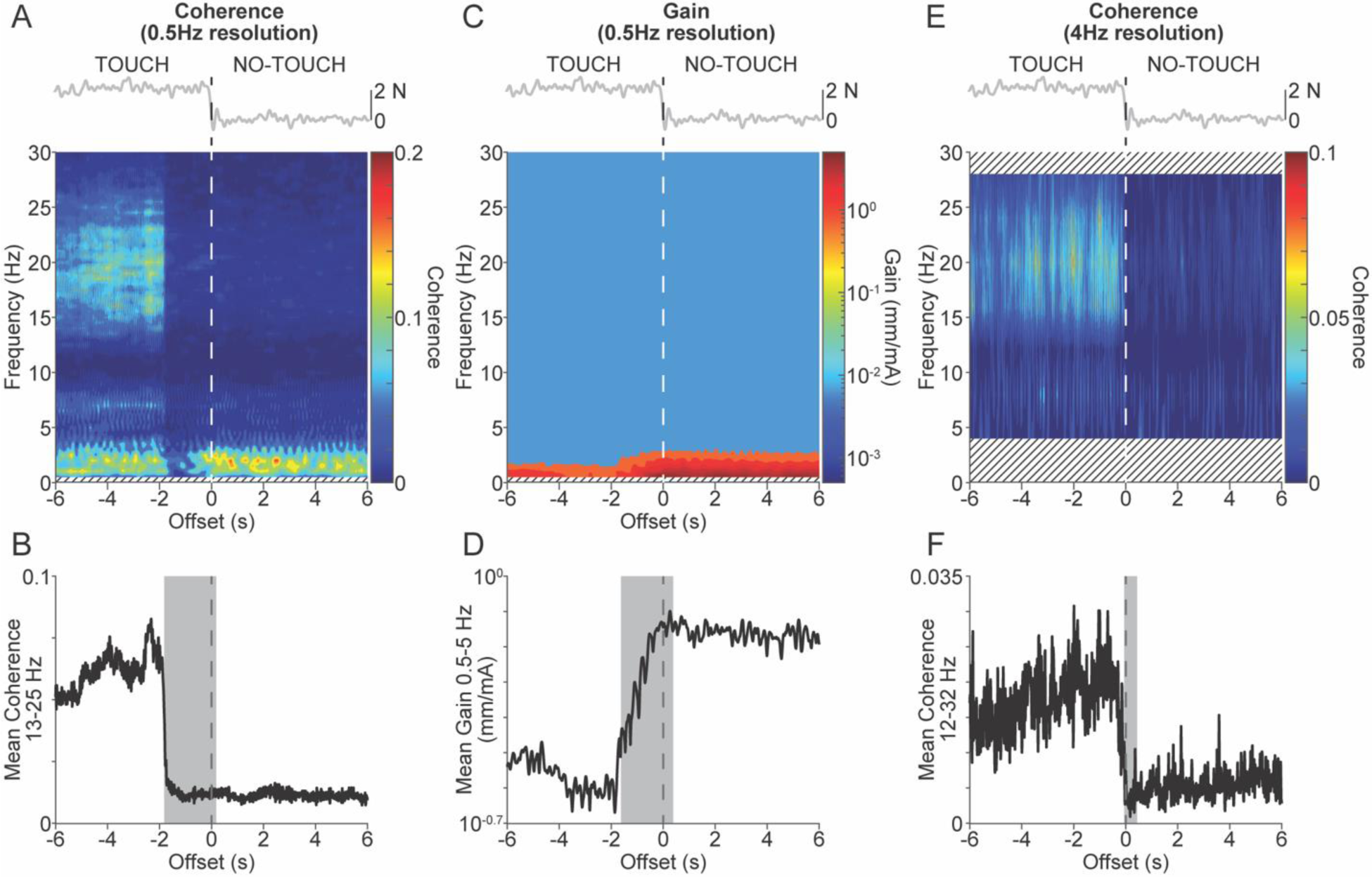
Time-Dependent Changes in EVS-CoP Coherence and Gain Before NO-TOUCH Transition. Panel A displays a 3D plot of pooled EVS-CoP coherence from TOUCH to NO-TOUCH. Significant coherence is represented in the heat map and non-significant coherence was set to zero (blue), while frequencies outside the chosen resolution were hashed to improve contrast. The y-axis represents frequency, and the x-axis time relative to the load cell breaking contact with the finger (time 0). Inset above shows load cell force from a representative participant at a TOUCH to NO-TOUCH transition, with time 0 indicated by a vertical dashed line. Panel B shows mean coherence (13-25 Hz) over the transition period, with the left edge of the grey box indicating the time where coherence dropped below the TOUCH level, and box width indicating the temporal resolution of the analysis (2s). Panel C illustrates pooled EVS-CoP gain changes on a log scale for NO-TOUCH transitions, with higher amplitudes in warmer colours. Panel D shows average time-dependent gain (0-5 Hz) with a grey box indicating the onset of the gain and window width. Panel E, using a 0.25 s window and 4 Hz resolution, depicts delayed coherence reductions before lifting in a 3D plot. Panel F shows mean coherence (12-32 Hz), with time where coherence is not different from NO-TOUCH level indicated by the left edge of the grey box, and temporal resolution denoted by box width.

Figure 5C shows time dependent gain, where low frequency gain (< 5 Hz) is denoted with the red colour. Gain was shown to increase 1.62 seconds before the load cell was dropped below the finger (Fig. 5D). These time dependent coherence and gain results indicate vestibulo-motor modulations occurring before transitioning to NO-TOUCH.

#### Switching to from NO-TOUCH to TOUCH

When conducting the time-dependent analysis of transitions to TOUCH, EVS-CoP coherence increased after touch occurred, while gain decreased before the load cell contacted the finger. Figure 6A shows time-dependent coherence, where the increased high frequency coherence (>13 Hz) after time 0 can be seen with the teal colour. Coherence began to increase 0.55 seconds after contact was made (Fig. 6B). Figure 6C shows time-dependent gain, where the decreased low frequency gain (< 5 Hz) before time 0 can be seen in the red colour. Gain was shown to begin to decrease 1.41 seconds before contact was made (Fig. 6D). These results indicate that vestibulo-motor modulations were different between removing versus adding TOUCH, where transitions to TOUCH led to delayed modulations.

**Figure 6:**
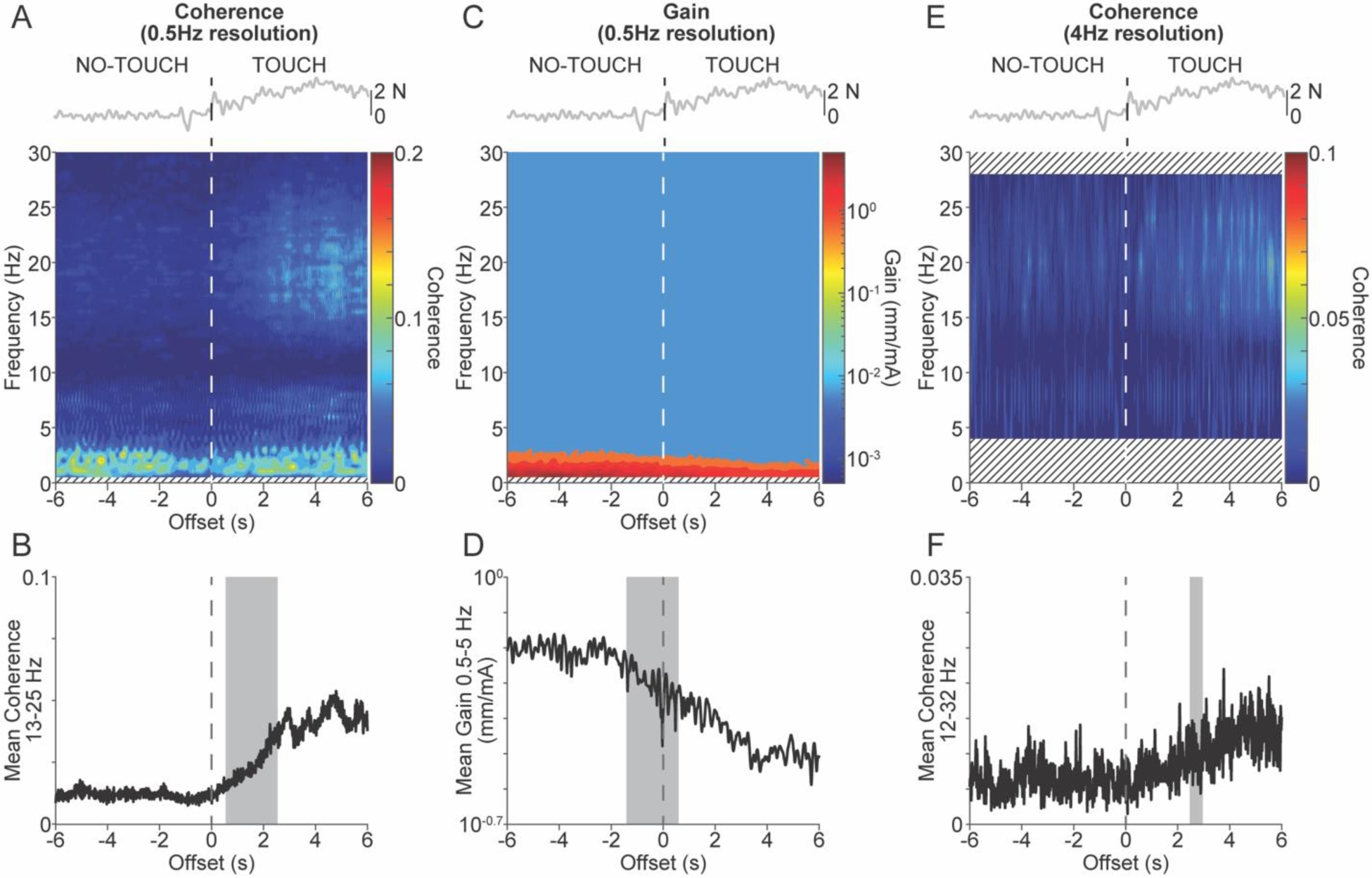
Time-dependent EVS-CoP coherence and gain changes occurred after transitions to TOUCH. All plots are the same as Figure 5 but show data from transitions to TOUCH (when finger contacts the load cell at time 0). Panel A displays a 3D plot of pooled EVS-CoP coherence from NO-TOUCH to TOUCH. Significant coherence is represented in the heat map and non-significant coherence was set to zero (blue), while frequencies outside the chosen resolution were hashed to improve contrast. The y-axis represents frequency, and the x-axis time relative to the load cell contacting the finger (time 0). Inset above shows load cell force from a representative participant at a NO-TOUCH to TOUCH transition, with time 0 indicated by a vertical dashed line. Panel B shows mean coherence (13-25 Hz) over the transition period, with the left edge of the grey box indicating the time where coherence increased above the NO-TOUCH level, and box width indicating the temporal resolution of the analysis (2s). Panel C illustrates pooled EVS-CoP gain changes on a log scale for TOUCH transitions, with higher amplitudes in warmer colours. Panel D shows average time-dependent gain (0-5 Hz) with a grey box indicating the onset of the gain and window width. Panel E, using a 0.25 s window and 4 Hz resolution, depicts delayed coherence increases after contacting in a 3D plot. Panel F shows mean coherence (12-32 Hz), with time where coherence is not different from TOUCH level indicated by the left edge of the grey box, and temporal resolution denoted by box width.

### Time-Dependent Signal Analysis

Since the apparent lead in vestibulo-motor coupling before switching to NO-TOUCH is highly unlikely, we conducted an additional post-hoc analysis using a 4 Hz resolution with 0.25 s windows to better resolve the timing of these adaptations. This revised analysis revealed that the detected onsets of coherence and gain changes may be influenced by the time windows selected.

For transitions to NO-TOUCH with the 0.25 s, or 4 Hz resolution, the pre-transition signal was significantly influenced by signal drift, leading to large standard deviations which were unsuitable for use in our onset detection algorithm. On visual inspection, an experimenter determined the decrease in coherence from the pre-transition mean occurred at approximately 0.28 s before the load cell was dropped below the finger (Fig. 5F). We then changed our approach to determine when the signal achieved its final state (as opposed to leaving the initial state). The coherence was not different from the post-transition state (within 2 standard deviations) at 0.06 s before the transition. For transitions to TOUCH, the pre-transition signal was not influenced by drift, and we used our detection algorithm. The increase in coherence began 2.46 seconds after the finger was contacted with the load cell (Fig. 6F).

### EVS-Segmental Linear Acceleration Relationship

Segmental kinematic analyses revealed that there was a trend towards greater EVS-acceleration coherence during NO-TOUCH at lower frequencies for all segments, and greater EVS-acceleration coherence during TOUCH at higher frequencies for the ankle and head. At the head, NO-TOUCH had significant EVS-acceleration coherence from 0.5-3, 4-8, 10.5-19.5 Hz while TOUCH was significant from 0.5-13.5, 14.5-22 Hz (Fig. 7A). NO-TOUCH had significantly greater coherence from 14-15 Hz while TOUCH had significantly greater coherence between 8.5-10 and 19.5-21 Hz (Fig. 7B). At the sternum, NO-TOUCH had significant coherence from 0.5-4, 6-8, 15-16 Hz while TOUCH had significant coherence from 0.5-8.5 Hz (Fig. 7D). At the low back, NO-TOUCH was significant from 0.5-3.5, 5-10.5 while TOUCH was significant from 0.5-2.5 and 7-8 Hz (Fig. 7G). Coherence was not significantly different between conditions for EVS-low back and sternum acceleration, but both trend towards NO-TOUCH having greater coherence at low frequencies (Fig. 7E, H). At the wrist, NO-TOUCH had significant coherence from 0.5-2, 5.5-6.5 and 7.5-9.5 Hz while TOUCH was significant from 0.5-1.5 and 4-7 Hz (Fig. 7J). NO-TOUCH had significantly greater coherence between 1-2 Hz (Fig. 7K). At the ankle, NO-TOUCH had significant coherence from 0.5-3, 4.5-7.5 and 8.5-16 Hz while TOUCH was from 0.5-2.5 and 8.5-18 Hz (Fig. 7M). TOUCH was significantly greater from 14.5 to 16 Hz (Fig. 7N), meaning the ankle was trending toward the EVS-CoP TOUCH high frequency coherence pattern slightly. Overall, variation in the segment acceleration that could be explained by EVS was increased during NO-TOUCH at the extremely low frequencies. For the ankle, there was more variation in the segment acceleration that could be explained by EVS during TOUCH at middle-high frequencies.

**Figure 7:**
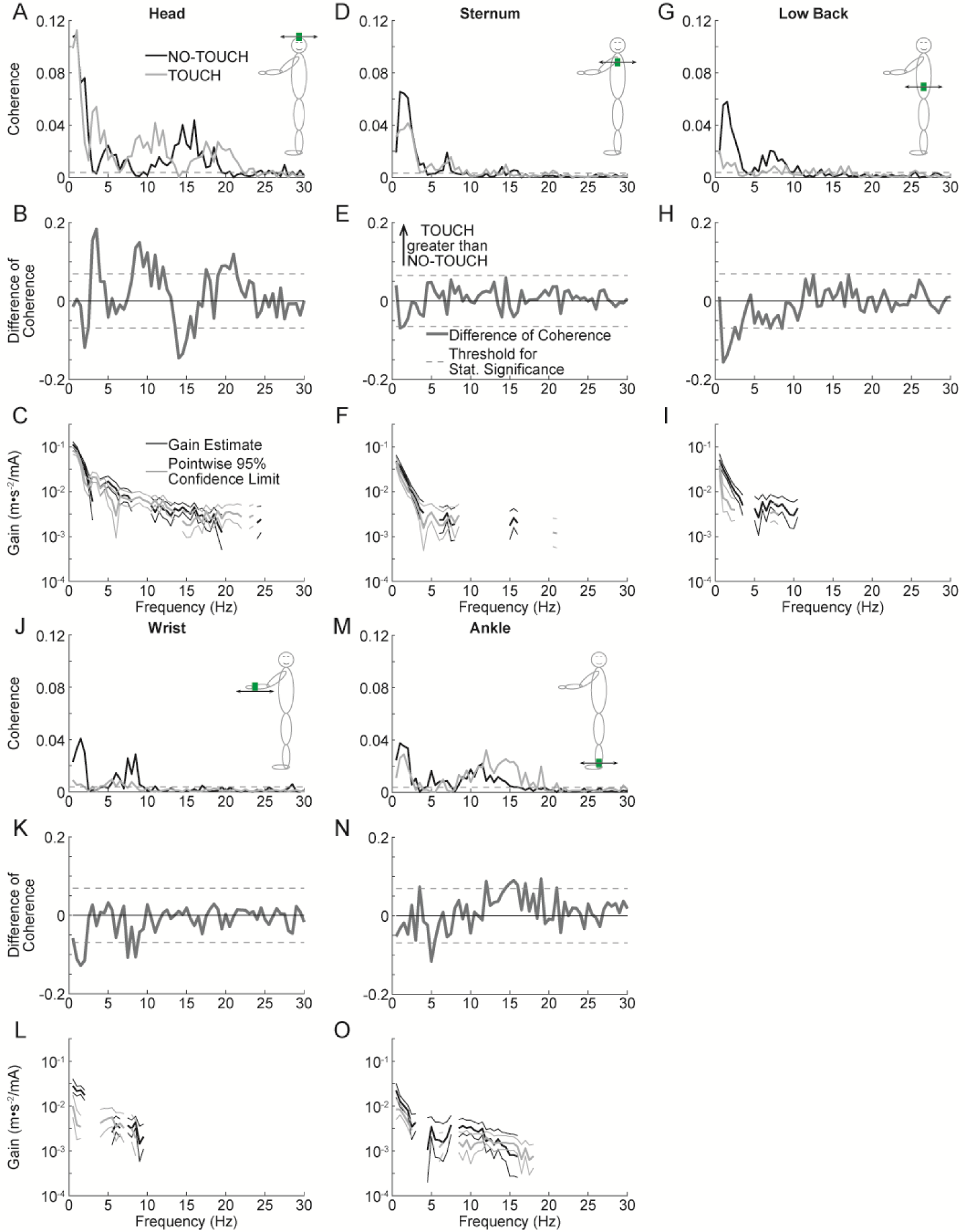
Coherence and Gain Differences Between TOUCH and NO-TOUCH in EVS-Segment Acceleration. Pooled EVS-acceleration coherence for head (A), sternum (D), low back (G), wrist (J), and ankle (M) for NO-TOUCH (black) and TOUCH (grey). A dashed horizontal line indicates the 95% confidence limit threshold for significant coherence. Accompanying images illustrate accelerometer placements and acceleration directions. Difference of coherence (black solid lines) are shown for each segment, with positive/negative 95% confidence limits (black dashed lines), where positive values indicate frequency ranges where TOUCH EVS-CoP coherence is greater than NO-TOUCH coherence. Lastly, EVS-CoP gain is shown for each segment on a log scale with pointwise 95% confidence limits (thin lines), excluding frequencies without significant coherence for clarity. Statistically significant differences in gain can be seen where 95% confidence limits do not overlap.

For head, sternum and ankle gain, there were no significant differences between NO-TOUCH and TOUCH, as there was no separation between gain estimates (solid lines) and their confidence limits (dotted lines) across conditions (Figs. 7C, F, O). EVS-linear acceleration gain was significantly greater in the NO-TOUCH condition between 1 and 2.5 Hz for the low back (Fig. 7I), and between 0.5 and 1.5 Hz for the wrist (Fig. 7L), matching the EVS-CoP gain results. Overall, there was more acceleration per unit of EVS at the low back and wrist during NO-TOUCH, while the acceleration at the head, sternum and ankle were similar across conditions.

## Discussion

This study systematically investigated the influence of minimal fingertip contact, or light touch, on the modulation of vestibulo-motor coordination in maintaining posture and balance. Specifically, we aimed to examine the complex interplay between newly introduced balance-relevant sensory cues from light touch and the intrinsic vestibulo-motor pathways, especially under conditions where vestibular information was discordant with gravitational cues. The main findings reveal that light touch attenuates the overall vestibular-evoked sway by reducing CoP-RMS and EVS-CoP gain but enhances the high-frequency coherence between EVS and CoP. This latter effect underscores an increased prevalence of EVS-induced variations in CoP at higher frequencies, a phenomenon not previously documented. Moreover, the temporal dynamics of coherence, exhibiting a decline prior to or shortly after the cessation of TOUCH, suggest a sophisticated mechanism within the CNS for adapting to changes in sensory cues at speed. When switching back to TOUCH, vestibulo-motor modulations take longer, suggesting potential asymmetrical responses to sensory cue changes. These findings extend our understanding of sensorimotor integration, demonstrating that the CNS not only dynamically reweights sensory inputs to optimize postural control but also incorporates salient cues like those from EVS, as they cannot entirely be discounted. Such insights contribute significantly to the broader discourse on multisensory feedback mechanisms in balance control and have potential implications for the design of therapeutic interventions and assistive technologies aimed at enhancing postural stability.

### Light Touch Reduces CoP Displacements

Our findings corroborate and extend upon previous investigations into the effects of light touch on the balance control system when measured as a whole. Notably, we observed a significant reduction in CoP displacement evidenced by decreases in both CoP RMS and EVS-CoP gain, ranging from 58-68% for gain during TOUCH, compared to NO-TOUCH. These data suggest light touch is both reducing spontaneous sway and reducing the amplitude of vestibular-evoked balance responses. This reduction is consistent with earlier studies (37, 40), who reported a decrease in CoP sway amplitude by 50 to 68% under similar experimental conditions. We restricted participants to under 2 N in the TOUCH condition because biomechanical models suggest mechanical stabilization of sway begins above 2 N (37). The degree to which participants gained a mechanical advantage was not assessed during TOUCH, and thus whether they were able to mechanically stabilize sway was not clear. It is unlikely that participants were mechanically stabilizing themselves significantly and these consistent observations across studies suggest that light touch serves as a reliable source of sensory information for balance control, potentially by enhancing the sensorimotor representation of the body’s position relative to gravity. Such a mechanism would allow the CNS to effectively counteract sway induced by potent vestibular perturbations such as those caused by EVS.

### Novel Insights: Light Touch Increases High Frequency Vestibular Contributions to Balance and Postural Control

Contrary to our initial hypothesis, we found that the presence of light touch led to an increase in EVS-CoP coherence, particularly at higher frequencies (11-28.5 Hz), despite a decrease in EVS-CoP gain. We initially expected coherence and gain to change in the same direction, which is a pattern seen in previous vestibular balance control research (5, 8, 9, 27, 30, 31). For example, Horslen and colleagues (5) found significantly increased coherence from 5.5-17.7 Hz and increased gain across all frequencies during increased height-induced postural threat. This divergence from our expectations and previous research highlights a unique modulation of vestibular processing by light touch. Specifically, light touch may influence vestibular information processing beyond merely affecting postural sway, indicating a novel aspect of sensorimotor integration.

The physiological significance of changes in high frequency EVS-CoP coherence was not immediately clear since most CoP power during unperturbed standing tends to occur under 1 Hz (49). However, the frequencies where EVS-CoP coherence were different between NO-TOUCH and TOUCH fall within the realm of natural vestibular stimulation, since people can experience vestibular inputs up to 20 Hz in situations like walking, stair climbing, running, jumping, sports, and riding the bus (50). Furthermore, leg and trunk muscles show responses to EVS up to 25 Hz during standing posture (51), postural transitions (31), and gait cycle (29, 46), while neck muscles show responses up to 150 Hz (52) independent of task demands (25, 53). While our experimental task had participants stand as still as possible, the vestibular system can operate at higher frequencies during dynamic balance situations to affect control of limb or segmental postural control and/or balance stabilization (29, 31, 46, 51, 52). Therefore, the observed changes in high frequency vestibulo-motor responses may reflect changes in vestibulo-motor coupling relevant to other balance or postural control scenarios. This understanding invites further exploration into how the CNS integrates and prioritizes sensory inputs during dynamic balance tasks, where high frequency vestibular inputs and responses are more prevalent. Future research could investigate how light touch influences sensory weighting and integration in more dynamic scenarios using frequency and amplitude matched-stochastic platform movements, simulating real-life dynamic conditions with equivalent vestibular stimulation. Such studies would provide insight into the adaptability of sensory weighting mechanisms under varying dynamic conditions, offering a richer understanding of balance maintenance.

### Temporal Dynamics of Sensory Reweighting

The timing of changes in EVS-CoP coherence and gain exhibited distinct patterns depending on whether participants were transitioning from TOUCH to NO-TOUCH or vice versa. Specifically, we detected a decrease in EVS-CoP coherence and increase in gain before the transition from TOUCH to NO-TOUCH. Conversely, transitioning from NO-TOUCH to TOUCH resulted in delays to increasing coherence while decreasing gain, with increased coherence occurring after the transition.

The decrease in high-frequency coherence before the unpredictable, and externally imposed transition from TOUCH to NO-TOUCH necessitated further investigation. While initial review of these data suggests anticipatory modulation of vestibulo-motor coupling and gain almost 2 s before the transition, there are several reasons related to experimental design and methodology to temper this conclusion. First, despite the intended unpredictability of transitions, there is a possibility participants anticipated changes, influenced by environmental cues or the mechanical sensation of load cell movement. Second, our time-dependent coherence and gain time estimates reflect the beginning of the analysis bin where a change was detected, meaning it could have occurred at that time or at any other time within the bin width. In our initial analyses, we used a 2 s bin-width to enable frequency resolution down to 0.5 Hz, where CoP power would be expected (49). We detected a decrease in high-frequency coherence from TOUCH to NO-TOUCH -1.82s before this transition, but this bin spans the transition event (-1.82s before to +0.18s after transition). As such, this pre-transition latency does not preclude a post-transition adaptation to vestibulo-motor coupling up to, and including, 0.18 s after the TOUCH to NO-TOUCH transition event. This window of uncertainty is illustrated with gray bands in figure 5. We subsequently reanalyzed these transitions using a 4 Hz frequency resolution (0.25 s bin width), which was just low enough to characterize the high-frequency range where changes in coherence between light touch conditions were observed. While signal drift precluded use of our 2 SD detection threshold to determine the onset of the decrease in coherence, visual inspection of these data suggests the change in coherence began –0.28 s before the TOUCH to NO-TOUCH transition (bin ending 0.03 s before). We could determine that coherence fully achieved the NO-TOUCH level –0.06 s before the transition (bin width representing –0.06 s before to 0.19 s after transition) by using a 2 SD threshold from the post-transition mean. As such, these data suggest the CNS is capable of adjusting vestibulo-motor coupling no later than 0.19 s after removal of light touch, and possibly in anticipation of removal of light touch.

Transitioning from NO-TOUCH to TOUCH resulted in a much later increase in coherence and decrease in gain, starting +0.55 s after (coherence) and –1.41 s before (gain) the introduction of light touch. Much like the latency of changes in coupling related to transition from TOUCH to NO-TOUCH, latencies related to the NO-TOUCH to TOUCH transition might be later than implied by our latency measures. As above, the algorithm-detected latencies reflect the beginning of an analysis bin where coherence (or gain) is different from the preceding bins. A +0.55 s latency might reflect a change in coherence and gain occurring up to +2.55 s after introduction of light touch. Reducing the bin width to 0.25 s to better resolve the latency of changes in high-frequency coherence led to a +2.46s latency (bin ending +2.71s). Combined, these data suggest high-frequency coherence is increasing in a delayed manner during transitions to TOUCH in comparison to transitions to NO-TOUCH. This nuanced pattern indicates that the CNS’s adaptation to sensory input changes may operate on a spectrum from anticipatory to reactive adjustments.

### Segmental Contributions to Balance or Postural Control

In Experiment 2, the inclusion of segmental kinematic analyses alongside CoP measurements allowed for a nuanced dissection of the vestibulo-motor pathways and their respective contributions to balance and posture control. This was motivated by the recognized variability in muscle involvement in response to EVS, which varies based on their role in balance and posture maintenance (54, 55), and differential modulation of vestibular-evoked responses across body segments (75). By examining head acceleration, we aimed to assess vestibulocollic reflex modulation, with additional measures at the sternum, low back, dominant arm, and ankles providing insights into whole-body sway, arm control via vestibulospinal modulation, and balance actions at the effectors closest to CoP generation.

Here, our analyses suggest that EVS activated vestibulocollic pathways to stabilize the head in both NO-TOUCH and TOUCH conditions, evidenced by the presence of EVS-head acceleration coherence across both conditions (Fig. 7A; NO-TOUCH: 0.5-3, 4-8, 10.5-19.5 Hz; TOUCH: 0.5-13.5, 14.5-22 Hz). This finding suggests an engagement of vestibulocollic reflexes within the high frequency range where changes in EVS-CoP coherence were also observed, however, it is unlikely that these reflexes were the primary drivers of the increased EVS-CoP coherence observed in the TOUCH condition. This inference is supported by the observation that both TOUCH and NO-TOUCH conditions exhibited greater coherence across distinct frequency bands (Fig. 7B; NO-TOUCH greater 14-15 Hz, TOUCH greater 8.5-10, 19.5-21 Hz). Despite this, we cannot completely rule out the role of vestibulocollic reflexes in contributing to the differences in EVS-CoP coherence across conditions, as there may be other neuro-mechanical mechanisms enhancing the transmission of head accelerations through the body to affect CoP during TOUCH. Nonetheless, the coherence and gain profiles for sternum and low back accelerations indicate minimal high-frequency action, suggesting these segments are unlikely conduits for transmitting coherent head accelerations to the feet in the TOUCH condition.

Arm stabilization emerged as a potential mediator of high-frequency EVS-CoP coherence, given the precise postural control required to maintain contact or proximity to the load cell in both TOUCH and NO-TOUCH conditions, and the known role of vestibular control for the arm during reaching tasks (56, 57). The tasks necessitated fingertip spatial positioning: in NO-TOUCH, participants hovered above the load cell without feedback; in TOUCH, participants maintained contact with the load cell (Experiment 1: 1 cm diameter; Experiment 2: 6 cm^2^) at a precise force level (1-2 N). This spatial motor control component, particularly pronounced during TOUCH with its heightened precision requirements and feedback on task performance, could theoretically influence vestibulo-motor coherence and gain. The CNS could increase vestibular weighting in the TOUCH condition to stabilize finger and hand ego- and allocentric posture in response to EVS-evoked vestibular perturbations. Yet, despite the presence of vestibular control over wrist accelerations in NO-TOUCH conditions, minimal EVS-wrist acceleration coherence was observed during TOUCH (Fig. 7J). This suggests that the increased precision in postural control did not significantly contribute to the EVS-CoP coherence. Therefore, the high-frequency EVS-CoP coherence observed during TOUCH is likely not attributable to vestibular control of arm stabilization and posture.

Our analyses indicate that the shank musculature may be actively responding to high frequency EVS, potentially driving the observed changes in EVS-CoP coherence in the TOUCH condition. Given the shank’s pivotal role at the end of the kinetic chain linking the body to the foot, its significant function in AP direction balance control, and the established lateral vestibulospinal connections to the shank musculature likely activated by EVS (25, 55), significant increases in EVS-ankle acceleration coherence within the TOUCH condition compared to NO-TOUCH (Fig. 7M; 14.5-16 Hz) were notable. Although aligned with the directional changes in EVS-CoP coherence, these effects were not as pronounced. However, since proximal segments (sternum and low back) do not show high frequency coherence to EVS, we suggest the biomechanical effects driving the high-frequency CoP response during TOUCH may originate in the shank. This is a hypothesis that remains to be directly tested due to the absence of muscle activity measurements in our study. Future investigations utilizing EMG could elucidate the vestibulo-muscular responses implied by our acceleration data, providing a clearer understanding of the underlying neuromechanical mechanisms.

### Mechanisms Underlying Dynamics of Vestibular and Cutaneous Integration

Our study posits that sensory reweighting within the CNS encompasses both anticipatory and reactionary adjustments, influenced by the introduction or removal of tactile cues while maintaining balance and posture. Previous research has identified patterns of anticipatory vestibulo-motor coupling adjustments during locomotion initiation or cessation (31) For example, Luu and colleagues (8) found a decrease in coherence 150-200 ms after participants were externally stabilized while balancing, and Rasman and colleagues (9) found coherence decreased 1.5 s following the introduction of sensorimotor delays. Analogous behaviour is observed in the cutaneous system, where removal of light touch prompts an immediate, albeit temporary, destabilization of sway, whereas the addition of touch results in a more gradual stabilization of postural sway (38). It takes about 1 s to destabilize sway during the withdrawal of light touch while it takes 0.7-2 s for the onset of CoP to reduce after removal of touch, and 1-3 s to reach a steady state (38).This dynamic response suggests a CNS mechanism that rapidly compensates for the loss of stabilizing input, potentially engaging proprioceptive feedback from leg muscles to refine sensory information critical for balance maintenance.

The differential response times for sensory reweighting, prompted by changes in tactile input, likely reflect the CNS’s evaluation of the threat level to balance or postural stability posed by these changes. Echoing findings from Jeka and colleagues (33) who found that visual gain adapts more rapidly to the loss than the gain of stabilizing information, we propose a mechanism where the CNS swiftly adjusts to the loss of tactile cues, given the immediate threat to stability. Further support for this comes from the work of Sozzi and colleagues (38) who suggested that the CNS rapidly detects the loss of the stabilizing input, and oscillating sway increases which engages proprioceptors in the leg muscles to gain more sensory information. The rapid adaptation following a loss of sensory information—a direct threat to balance and posture—suggests a crucial, possibly anticipatory, CNS response to mitigate postural instability or fall risk. Similarly, Phanthanourak and colleagues (58) reported that increased postural threat due to support surface translations leads to larger anticipatory postural adjustments to mitigate fall risk. Conversely, the incorporation of new sensory data during TOUCH presents a less immediate threat, allowing a more deliberative or cautious CNS evaluation and adjustment period. This nuanced response time might reflect an evolutionary adaptation, prioritizing the maintenance of balance and posture and minimization of fall risk above the rapid integration of novel, potentially non-essential sensory cues. This hypothesis points to the sophisticated balance and posture maintenance strategies employed by the CNS, highlighting the selective and adaptive nature of sensory reweighting in response to dynamic environmental changes and the perceived immediacy of balance and posture threats.

We propose that the cutaneous system is well-suited to effecting dynamic changes in vestibulo-motor coupling at high frequencies (13-30 Hz). Fast adapting type I (FAI) cutaneous receptors, which are dense in the fingertips, preferentially fire in midrange frequencies (10–50 Hz; (59, 60)), encapsulating the profile of significantly different coherence observed here (13-30 Hz). There are many neural mechanisms through which cutaneous inputs might influence high frequency vestibular-evoked balance responses, including spinal (61, 62) and supra-spinal cutaneous reflex circuits that share similar end targets as vestibulospinal circuits (63–65), as well as integration with vestibular sensory inputs to modulate vestibulospinal function and/or multi-sensory balance and posture control at the brainstem (66), cerebellum (67), or cortical levels (22, 68–70).

During TOUCH, the EVS-driven vestibular signal is perceived as involuntary rotation (71), causing changing pressure between the finger and the light touch surface and counteracting balance responses (72). It has been shown that displacement of a light touch reference can lead to short-latency spinal reflexes in ankle muscles consistent with a balance reaction (62). Reflexes via the reticulospinal tract, which innervate large groups of muscles in synergistic patterns (73) and is involved in restoring postural control (63–65), could also be triggered by interactions between the finger and surface. Since these potential cutaneous-driven balance reactions are counteracting the movement caused by EVS, CoP may appear to cohere more with EVS during TOUCH at high frequencies, where FAI receptors encode touch information.

Observed changes in vestibulo-motor coupling may also be driven by modulation of vestibular reflexes. In this case, the vestibular nuclei would likely be involved, where EVS-driven reflexes originate multisensory integration with sensory input like vestibular and somatosensory cues occurs (25, 66, 74, 75). Specifically, the inferior vestibular nuclei receive input from the vestibular afferents and vermis of the cerebellum and is thought to be a site where integration between vestibular and cutaneous input can occur (76). Descending input to the lateral vestibulospinal tract controlling low frequency responses, where postural sway occurs, may have been inhibited when light touch was present, reflected in the EVS-CoP gain reductions. High frequency light touch information from FAIs may modulate high frequency vestibular responses by increasing the likeliness that excitatory neurons can be brought to threshold by EVS, reflected by the increased high frequency EVS-CoP coherence.

Light touch is also known to induce activation in several multisensory cortex regions which could influence vestibular reflex responses, including the posterior parietal cortex (68, 77), vestibular cortex (70), somatosensory cortex (69), insula, superior temporal gyrus, supramarginal gyrus, striatum, amygdala, and prefrontal cortex (78). Additionally, the cerebellum receives multisensory input and has inhibitory control over the vestibular nuclei through the vestibulocerebellum network (67, 79). The likeliness of cortex structures or the cerebellum playing a role in vestibular reflex modulation increases when considering the longer re-weighting period during the transition to TOUCH ((75) refer to Fig. 6). Vestibulo-motor adaptations driven by cutaneous reflex circuits or cutaneous-vestibular interactions within the vestibular nuclei, cerebellum, or cortex are all speculations, as we did not record output from these CNS structures, and this was not the goal of the study.

### Concluding Remarks

In conclusion, the results from this study demonstrate how vestibulo-motor control of balance and posture is altered by light touch, and the time it takes to do so. Light touch decreased EVS-CoP gain while increasing coherence, meaning there was less CoP displacement, but more CoP variation attributed to EVS. Our unique findings show that light touch can re-weight vestibular-evoked responses by reducing their size but also increasing high frequency vestibular contributions for the remaining sway. This suggests that the CNS can use novel sensory inputs to alter balance and posture but cannot completely ignore a salient balance/postural cue. Time dependant analyses revealed that vestibular modulations occurred before removing light touch, but after addition, showing that there can be response timing asymmetries between introduction versus removal of sensory information. The loss of sensory information may be more destabilizing, requiring faster adjustments, while it may take more time for the CNS to determine the relative threat to balance and posture light touch imposes upon introduction.

This study aimed to provide insight into the underlying mechanisms for the sensorimotor control of balance and posture, but cautious interpretation is still needed when generalizing results to other specific balance-related situations. The complexity of multisensory integration to maintain balance and posture has been demonstrated, where the ability of one sensory modality to modulate another by the CNS is not always easy to predict. Providing light touch feedback may be a useful strategy to enhance vestibular cues, especially in dynamic situations where vestibular feedback has a larger high frequency component. Light touch could be used during therapeutic interventions or for assistive technologies aimed at enhancing vestibular deficits or postural stability. This study also motivates further research focusing on sensory interactions in balance or postural control.

## Data Availability

All data and code for this article are provided in an Open Science Framework (OSF) repository, at https://osf.io/gy98m/. We report all measured variables and all analyses we conducted.

## Acknowledgements

We thank Jeff Rice for helping to construct the load cell arms of the light touch apparatus, Lukas Forbes for assisting with collection and processing of accelerometer data, Ben Cornish for late draft review, as well as Bill McIlroy and Andrew Laing for scientific discussion.

## Disclosures and Disclaimers

The authors have no disclosures or disclaimers to declare.

## Author Contributions

BCH devised the main concept of the project and the theoretical framework. BCH and MHG conceived the original experiments. MHG and BCH carried out the experiments and analyzed the data. MHG took the lead in writing the manuscript. BCH and MBC secured funding for the project and oversaw overall direction and planning. All authors provided critical feedback and helped shape the research, analysis, and manuscript.

