## Supplementary Figures for "Dynamics of sensorimotor reweighting: How light touch alters vestibular-evoked balance responses"

### Supplementary Materials

#### EVS-ML CoP Relationship

Light touch minimally affected the relationship between EVS and ML-CoP, leading to a focus on the results from AP-CoP. There was significant EVS–ML CoP within-conditions coherence in experiment 1 pooled data from 0.5 Hz to 19 Hz for NO-TOUCH and from 0.5-17.5, 18.5-19.5, 20.5-24.5 and 25.5-27 Hz for TOUCH (SFig. 1A). Significant coherence was determined when calculated coherence exceeded a 95% confidence limit at 0.0018 $R^2$. Additionally, EVS–ML CoP coherence was significantly greater with TOUCH compared to NO-TOUCH for a small range, from 0.5-1.5 Hz (SFig. 1B). Response size amplitude to EVS was similar for NO-TOUCH and TOUCH, where gain was significantly lower in the TOUCH compared to NO-TOUCH condition from 6.5 Hz to 8 Hz (SFig. 1C). EVS-CoP gain decreased logarithmically between 0.5- 10 Hz in both the NO-TOUCH and TOUCH conditions.

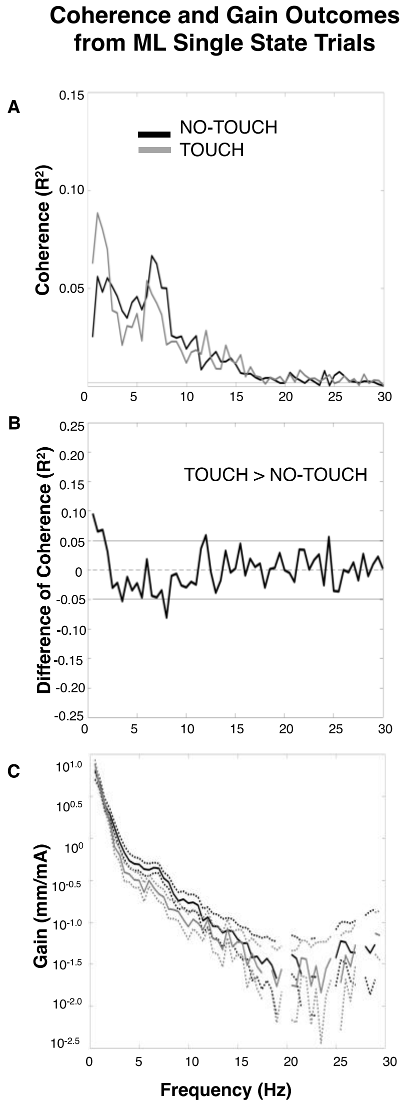

**Supplementary Figure 1: TOUCH Impact on EVS-ML CoP Coherence and Gain**

Panel A presents EVS-ML-CoP coherence across 16 participants, with NO-TOUCH in black and TOUCH in grey. A dashed horizontal line indicates significant coherence. Panel B shows difference of coherence (black solid lines) with positive/negative 95% confidence limits (black dashed lines), where positive values indicate frequency ranges where TOUCH EVS-ML-CoP coherence is greater than NO-TOUCH coherence. Panel C presents EVS-CoP gain on a log scale with pointwise 95% confidence limits (thin, dashed lines), excluding frequencies without significant coherence for clarity. Statistically significant differences in gain can be seen where 95% confidence limits do not overlap.

#### CoP Spectral Power

Reductions in CoP spectra were found from NO-TOUCH to TOUCH in both experiments. CoP amplitude normalization was conducted on all participants and conditions to ensure that coherence was being driven by the cross spectrum between EVS and CoP, and not these changes in CoP auto-spectra. SFig. 2 Panels A and C show CoP amplitude over 60 s for a representative participant from experiment 1 and 2, respectively, for NO-TOUCH and TOUCH conditions, where NO-TOUCH amplitude was greater. Amplitudes were normalized in Panels B and D, causing amplitudes to be more similar across conditions. Panel E and G shows how the pooled data CoP auto-spectrum was reduced in non-normalized TOUCH for experiment 1 and 2, respectively. After normalization, panels F and J shows that CoP auto-spectra became more similar for both experiments. Difference of spectra tests in Panels I and K revealed that NO-TOUCH had significantly greater spectra across 0-30 Hz for both experiments. When CoP amplitude was normalized, the range and amplitude of where NO-TOUCH had significantly greater spectra was reduced. NO-TOUCH had greater spectra between 2.5-8, 9.5-19.5 and 20.5-30 Hz in experiment 1 group-wide data (SFig. 2D) and between 2-15.5 and 25-30 Hz in experiment 2 group-wide data (SFig. 2J).

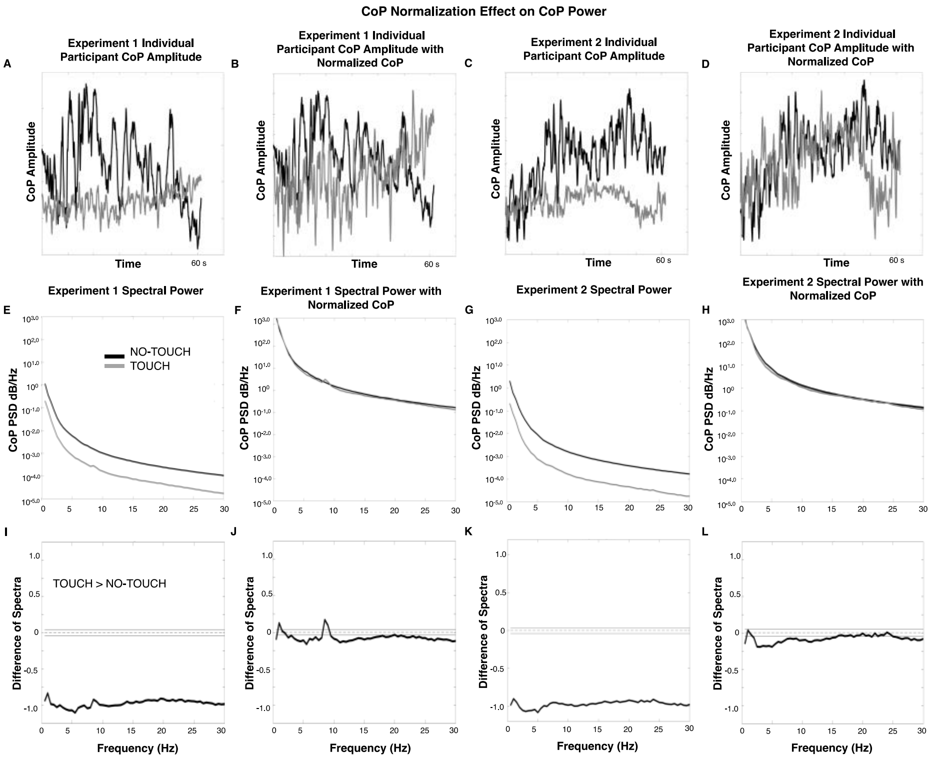

**Supplementary Figure 2: CoP Spectra Reduction During TOUCH Eliminated by Amplitude Normalization.**

Panels A and C display 60 s CoP amplitudes for a representative participant in experiments 1 and 2 under NO-TOUCH (black) and TOUCH (grey). Panels B and D show these series post-amplitude normalization. Panels E and G present pooled auto-spectrum data for both conditions and experiments, with F and H depicting normalized data. Panels I and K illustrate spectral differences for non-normalized CoP (black solid lines) with positive/negative 95% confidence intervals (dashed black lines), where positive values indicate frequency ranges where TOUCH spectra are greater than NO-TOUCH spectra. Panels J and L show the difference in spectra after normalization.

#### EVS-CoP Relationship During Switching Trials

The data from the switching trials exhibited patterns of results that were like the single state trials. This confirms the relationship between EVS and CoP remained similar despite the transitions between conditions. There were 380 transitions to NO-TOUCH used in this sample, with 8 seconds on either side of the transition included. Data were concatenated in order of collection across all participants (n = 10) for group-wide analyses to yield a single 1520-bin array of EVS and CoP data for each condition, where each participant contributed an equal amount of data to the analyses. There was significant EVS–CoP within-conditions coherence from 0.5-9.5, 12.5-23 and 24-27.5 Hz for NO-TOUCH and from 0.5-9.5 and 11.5-30 30 Hz for TOUCH (SFig. 3A). Significant coherence was determined when calculated coherence exceeded a 95% confidence limit at 0.00197 $R^2$. EVS–CoP coherence was significantly greater with TOUCH compared to NO-TOUCH from 6.5-7.5, 13-27 and 28-29 Hz (SFig. 3C). The gain between EVS and CoP was significantly decreased with light touch available from 0.5 Hz to 6 Hz, shown in panel E. EVS-CoP gain decreased logarithmically between 0.5 Hz and 10 Hz in both the NO-TOUCH and TOUCH conditions. This data means that time-dependent results should show similar patterns before and after the transition periods.

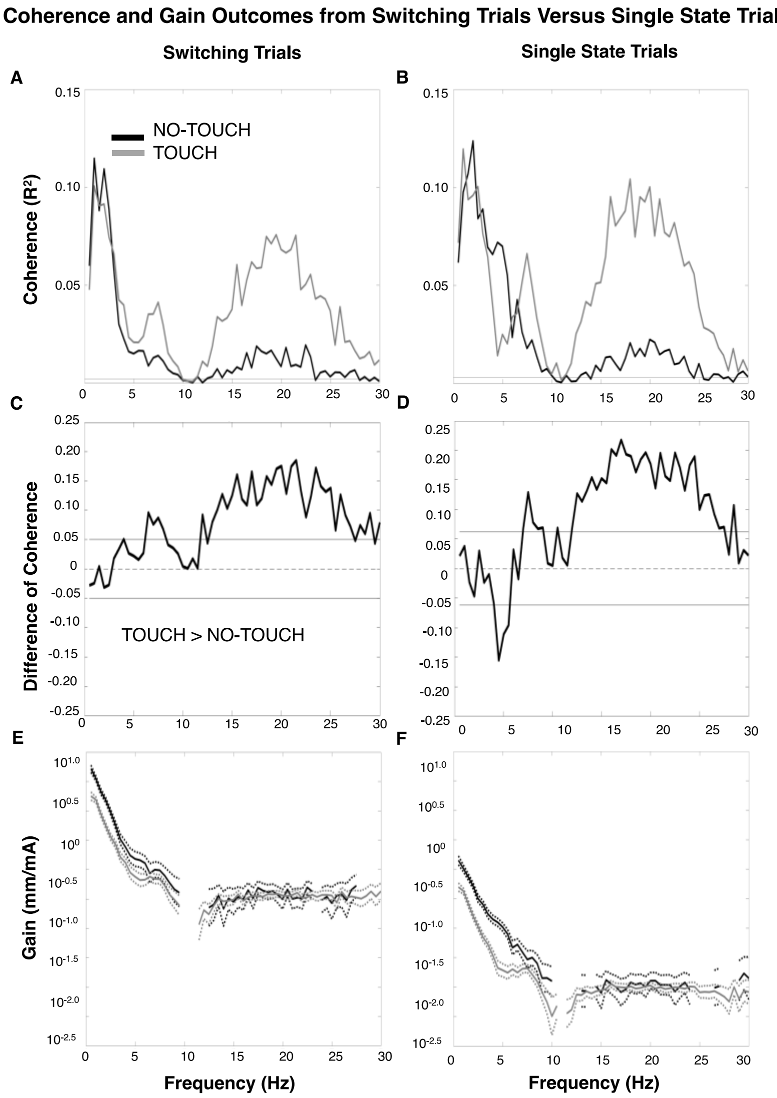


**Supplementary Figure 3: Consistent TOUCH Effects on EVS-CoP Coherence and Gain in Switching vs. Single State Trials.**

Panels A and B depict EVS-CoP coherence for experiment 2's single state (A) and switching trials (B), with NO-TOUCH in black and TOUCH in grey. A dashed horizontal line indicates significant coherence. Panels C and D show difference of coherence (black solid lines) with positive/negative 95% confidence limits (black dashed lines), where positive values indicate frequency ranges where TOUCH EVS-CoP coherence is greater than NO-TOUCH coherence. Panels E and F show EVS-CoP gain on a log scale with pointwise 95% confidence limits (thin, dashed lines), excluding frequencies without significant coherence for clarity. Statistically significant differences in gain can be seen where 95% confidence limits do not overlap.
